# The cGAS-STING pathway affects vertebral bone but does not promote intervertebral disc cell senescence or degeneration

**DOI:** 10.1101/2022.03.17.484614

**Authors:** Olivia K. Ottone, Cheeho Kim, John. A. Collins, Makarand V. Risbud

**Author notes:** **Correspondence:** Makarand V. Risbud, Ph.D., Department of Orthopaedic Surgery, Sidney Kimmel Medical College, 1025 Walnut Street, Suite 501 College Bldg., Thomas Jefferson University, Philadelphia, PA 19107.

## Abstract

The DNA-sensing cGAS-STING pathway promotes the senescence-associated secretory phenotype (SASP) and mediates type-I interferon inflammatory responses to foreign viral and bacterial DNA as well as self-DNA. Studies of the intervertebral disc in humans and mice demonstrate associations between aging, increased cell senescence, and disc degeneration. Herein we assessed the role of STING in SASP promotion in STING gain- (N153S) and loss-of-function mouse models. N153S mice evidenced elevated circulating levels of proinflammatory markers including IL-1β, IL-6, and TNF-α and exhibited a mild trabecular and cortical bone phenotype in caudal vertebrae. Interestingly, despite systemic inflammation, the structural integrity of the disc and knee articular joint remained intact, and cells did not show a loss of their phenotype or elevated SASP. Transcriptomic analysis of N153S tissues demonstrated an upregulated immune response by disc cells, which did not closely resemble inflammatory changes in human tissues. Interestingly, STING^-/-^ mice also showed a mild vertebral bone phenotype, but the absence of STING did not improve the age-associated disc phenotype or reduce the abundance of SASP markers. Overall, the analyses of N153S and STING^-/-^ mice that the cGAS-STING pathway is not a major contributor to SASP induction and consequent disc aging and degeneration but may play a minor role in the maintenance of trabecular bone in the vertebrae. This work contributes to a growing body of work demonstrating that systemic inflammation is not a key driver of disc degeneration.

## Introduction

Intervertebral disc degeneration is widely accepted as a major risk factor for chronic low back pain and neck pain, two of the leading causes of years lived with disability worldwide (1,2). The pervasive nature of these pathologies drives the need to understand the underlying mechanisms of major risk factors like disc degeneration. The disc constitutes the largest avascular, and therefore hypoxic, tissue in the body, and cellular function is tightly governed by the transcription factor HIF-1α (3–7). The disc is comprised of three compartments: an inner, glycosaminoglycan (GAG)-rich nucleus pulposus (NP); a fibrocartilaginous annulus fibrosus (AF), consisting of concentric collagen-rich lamellae surrounding the NP; and inferior and superior cartilaginous endplates (EP), which anchor the disc to adjacent vertebrae and facilitate nutrient diffusion between the vascularized vertebrae and the avascular disc (8). During aging and degeneration, water-imbibing proteoglycans are lost in the NP, and cells undergo morphologic and phenotypic changes, contributing to fibrotic remodeling in the NP and AF compartments and a reduced ability to accommodate spinal loading (9,10).

A widely recognized risk factor for disc degeneration, a slow progressing pathology, is aging (11). An important contributing factor to age-associated degeneration in the disc and knee joint articular cartilage is cell senescence. In human tissues, the incidence of senescence increases as these tissues age and during degeneration, marked by characteristic SA-β-gal staining, telomere shortening, and elevated P16^INK4A^ expression (12–15). The Ercc1^-/Δ^ progeria mouse model demonstrates genotoxic stress-induced premature senescence and disc degeneration (16,17). Senescence is of particular interest in degenerative disorders because of the senescence-associated secretory phenotype (SASP) developed by senescent cells. Characteristic of SASP is the secretion of various inflammatory cytokines and proteases – including known markers of disc degeneration IL-6, IL-1, MCP-1, and MMP-1, 3 and 13 – which contribute to the catabolism within tissues and propagate SASP (18,19). Targeting of senescent disc cells with the senotherapeutic o-vanillin has shown efficacy in reducing SASP in vitro, and most recently, the combination senolytic drug treatment of Dasatinib and Quercetin reduced SASP and incidence of disc degeneration in vivo, in aged C57B6 mice (20,21). These studies strongly support the need to further explore and understand the underlying mechanisms which drive disc cell senescence.

One such mechanism which has been demonstrated to promote SASP is the cGAS-STING DNA-sensing pathway which mediates type-I interferon inflammatory responses to foreign, viral, and bacterial DNA as well as self-DNA (22). It has been shown that cellular senescence destabilizes the nuclear membrane, resulting in the release of cytosolic chromatin fragments and consequent activation of the cGAS-STING pathway (23,24). Notably, Sting-null mice show signs of attenuated SASP after exposure to ionizing irradiation, indicating the cGAS-STING pathway is a critical mediator of DNA damage-associated SASP (24). STING has also been implicated in mediating inflammatory processes in musculoskeletal tissues. Deletion of STING in DNase^+/-^; Ifnar^-/-^ double knockout (DKO) mice – an inflammatory model used to study rheumatoid arthritis – effectively ameliorated paw inflammation and abnormal bone accrual in the long bones and spleens of the DKO animals (25,26). Recently, a rat model of vertebral inflammation-induced intervertebral disc degeneration, where lipopolysaccharide (LPS) was introduced into vertebral defects, showed a degenerative phenotype marked by cartilaginous endplate defects, immune cell infiltration, and higher levels of cGAS, STING, TBK1, and IRF3 (27). Likewise, an acute injury model of disc herniation was used by Zhang et al. to show that inhibition of STING results in the partial rescue of a degenerative phenotype by modulating NLRP3 signaling (28).

Based on these relationships between disc degeneration, senescence, and STING, we first investigated if, in absence of a traumatic mechanical injury, activation of STING in the intervertebral disc would promote SASP and a degenerative phenotype and secondly whether the absence of STING would be protective to the intervertebral disc by delaying the SASP program and reducing the prevalence of the age-related degeneration in mice. This was achieved by evaluating the spines of mice heterozygous (N153S) for the Sting^1em1Jmin^ mutation that constitutively activates STING without altering STING expression levels and STING^-/-^ mice to assess the contribution of STING to SASP onset and intervertebral disc degeneration (24,29). Interestingly, neither model demonstrated significant changes in the disc phenotype, with analyses of N153S mice serving as additional support to an emerging concept that when the structural integrity of the disc remains intact, the NP compartment is isolated from systemic inflammation (30). Overall, the analyses of the vertebral columns of N153S and STING^-/-^ mice indicate that the cGAS-STING pathway is not a major contributor to SASP induction and disc degeneration but may play a small role in the maintenance of trabecular bone in the vertebrae.

## Materials and Methods

### Animals

Animals procedures were performed under approved protocols by the IACUC of either Thomas Jefferson University (WT, N153S) or the University of Pennsylvania (STING^+/+^, STING^-/-^). Wildtype (WT) and heterozygous B6J.B6N-Sting1^em1Jmin^ (Stock # 033543) knock-in mice with a N153S substitution in sting1 gene were obtained from the Jackson Laboratory and aged to 6 months of age. WT and N153S animals were collected at 6 months-of-age due to poor survival in N153S mice by one year, resulting from multiple pathologies including progressive lung and perivascular inflammation and T cell cytopenia (29,31). Tissues from female and male animals were evaluated for evidence of accelerated disc degeneration, osteoarthritis, and SASP. Spines of 16-18 month-old STING^+/+^ and STING^-/-^ mice on a C57BL/6 were provided by Dr. Shelley Berger from the University of Pennsylvania (24,32). For each genotype, tissues from female and male animals between 16 and 18 months of age were evaluated for evidence of delayed age- and senescence-related degeneration.

### Plasma collection and analysis

Blood from 6-month-old WT (n = 9) and N153S (n = 9) mice was collect immediately postmortem by intracardiac puncture using heparinized needles. Plasma was separated from red blood cells via centrifugation at 1500 rcf and 4°C for 15 minutes and stored at −80°C until the time of analysis. Cytokine concentrations were evaluated using the V-PLEX Mouse Cytokine 19-Plex Kit (Meso Scale Diagnostics, Rockville, MD) according to the manufacturer’s specifications.

### Micro-Computed Tomography (μCT) Analysis

μCT scans (Bruker Skyscan 1275; Bruker, Kontich, Belgium) were performed on spines from all genotypes (n=6-7 mice/genotype) fixed with 4% PFA. An aluminum filter was used, and all scans were conducted at 50 kV and 200 μA, with an exposure time of 85 ms, yielding a resolution of 15 mm. Three-dimensional image reconstructions were generated from lumbar (L3-L6: WT, N153S; L4-L6: STING^+/+^ STING^-/-^) and caudal (Ca5-Ca9: WT, N153S; Ca5-Ca7: STING^+/+^ STING^-/-^) scans in CTan (Bruker) and used for all subsequent analyses. Intervertebral disc height and vertebral length were measured and used to calculate the disc height index (DHI), as previously described (33,34). The 3-D microarchitecture of the trabecular bone was tabulated in a region of interest (ROI) defined by contouring the outer boundary of the trabeculae throughout the vertebral body. Resulting datasets were assessed for the following parameters: structure model index (SMI), bone volume fraction (BV/TV), trabecular number (Tb. N.), trabecular thickness (Tb. Th.), and trabecular separation (Tb. Sp.). The cortical bone was analyzed in two dimensions and assessed for bone volume (BV), cross sectional thickness, mean cross-sectional bone area, and bone perimeter. Mineral density was calculated in lumbar vertebrae and boney endplates using a standard curve created with a mineral density calibration phantom pair (0.25 g/cm3 calcium hydroxyapatite (CaHA), 0.75 g/cm3 CaHA).

The hindlimbs WT and N153S (n=2 limbs/animal, 3 animals/genotype, 6 limbs/genotype) were scanned using an aluminum filter at 70 kV and 142 μA, with an exposure time of 55 ms, yielding a resolution of 7 μm. 3-D analysis as described above was conducted on the knee joints, and BV/TV, Tb. Th., Tb. Sp., and the subchondral bone plate thickness (SCBP Th.) were assessed between genotypes.

### Biomechanical Analysis

Caudal vertebrae (Ca2-4) from 6-month-old WT and N153S mice (n=12 animals/genotype) were isolated and stored in PBS-soaked gauze at −80°C before use. Samples underwent two freez-thaw cycles and were scanned using μCT prior to testing. Each vertebra was individually potted into a 2-mm plastic ring mold using an acrylic resin (Ortho-Jet, Patterson Dental, Saint Paul, MN), and mechanical loading was applied using a material testing system (TA Systems Electrofoce 3200 Series II). A 0.4-N compressive preload was applied, followed by a monotonic displacement ramp at 0.1 mm/s until failure. Force-displacement data were digitally captured at 25 Hz and converted to stress-strain using a custom GNU Octave script with μCT-based geometric measurements, as previously described (35,36).

### Histological Analysis

Spine segments used for calcified sections (WT and N153S: L1-L2, Ca1-Ca2) were fixed for 2 hours in 4% PFA in PBS, treated with 30% sucrose, OCT-embedded, and snap-frozen. For all genotypes, caudal and lumbar spines were dissected and immediately fixed in 4% PFA in PBS at 4°C for 48 hours. 18 days of decalcification in 20% EDTA at 4°C followed, and spines were then embedded in paraffin. Coronal sections of 7 μm were generated from caudal (Ca5-Ca9: WT, N153S; Ca3-Ca7: STING^+/+^ STING^-/-^) and lumbar (L3-LS1: WT, N153S; L1-S1: STING^+/+^ STING^-/-^) discs. Histoclear deparaffinization followed by graded ethanol rehydration preceded all staining protocols. Caudal (n=4 discs/mouse, 6-7 mice/genotype) and lumbar (n=4-6 discs/mouse, 6-7 mice/genotype) tissues were stained with Safranin O/Fast Green/Hematoxylin or Picrosirius red and imaged using a light microscope (AxioImager 2; Carl Zeiss Microscopy, Peabody, MA, USA) or a polarizing light microscope (Eclipse LV100 POL; Nikon, Tokyo, Japan). Imaging of Safranin O-stained tissues was performed using 5x/0.15 N-Achroplan and 20x/0,5 EC Plan-Neofluar (Carl Zeiss) objectives and Zen2^TM^ software (Carl Zeiss). Six blinded graders scored NP and AF compartments, using Modified Thompson Grading (37). Picrosirius red-stained tissues were imaged using 4x and 20x 10×/0.25 Pol/WD 7.0 (Nikon) objectives, and the areas occupied by green, yellow, or red pixels were determined using the NIS Elements Viewer software (Nikon).

WT and N153S hindlimbs (n=1 limb/animal, 3 animals/genotype, 3 limbs/genotype) were processed according to the aforementioned fixation and decalcification, protocols. 5 μm sections cut in the coronal plane underwent deparaffinized followed by graded ethanol rehydration. Midcoronal sections were stained with toluidine blue or H&E and imaged using a light microscope with 10x objectives (Motic BA300 POL; Motic, Schertz, TX, USA) and Jenoptik ProgRes Speed XT Core 5 camera with the ProgRes Capture Pro software (version 2.10.0.1) (Jenoptik Optical Systems). Histological scoring of cartilage damage, OA severity, osteophyte formation, and chondrocytes cell death was conducted using the ACS scoring system, as previously detailed (38,39).

### TRAP and TNAP Staining

Frozen calcified sections of WT and N153S discs (L1-L2, Ca1-Ca2) were cut at 10 μm and secured to slides using cryofilm 2C(10) to maintain the morphology of the mineralized sections (40). The taped sections were glued to microscope slides using chitosan adhesive and rehydrated prior to imaging. Sections were also stained to detect activity of tartrate-resistant acid phosphatase (TRAP) (1:80, Invitrogen, E6601A) and tissue non-specific alkaline phosphatase (TNAP) (Vector Laboratories, SK5300).

### TUNEL assay and cell number quantification

TUNEL assay was performed on N153S and WT lumbar disc tissue sections using an in situ cell death detection kit (Roche Diagnostic). Briefly, sections were deparaffinized and permeabilized with Proteinase K (20 mg/ mL) for 15 min at room temperature. The TUNEL assay was then carried out per the manufacturer’s protocol. The sections were washed and mounted with Prolong Gold Antifade Mountant with DAPI (Thermo Fisher Scientific, P36934). All mounted slides were imaged with an Axio Imager 2 microscope using 5 x /0.15 N-Achroplan or 10 x /0.3 EC Plan-Neofluar or 20 x /0.5 EC Plan-Neofluar objectives (Carl Zeiss) objectives, X-CiteÒ120Q Excitation Light Source (Excelitas), AxioCam MRm camera (Carl Zeiss), and Zen2TMsoftware (Carl Zeiss). TUNEL-positive cells and DAPI-positive cells were analyzed respectively to assess cell death and cell number in disc compartments

### Immunohistochemical Analysis

All immunohistochemical stains were conducted on mid-coronal lumbar sections of 7μm. Sections were deparaffinized and rehydrated, as described under Histological Analysis. Antibody-specific antigen retrieval was conducted by way of incubation in either a hot citrate buffer for 20 minutes, proteinase K treatment for 10 minutes at room temperature, or chondroitinase ABC for 30 minutes at 37°C. Sections were then blocked in 5% normal serum (Thermo Fisher Scientific, 10,000C) in PBS-T (0.4% Triton X-100 in PBS), and incubated with primary antibodies against Aggrecan (1:50; Millipore; AB1031), CA3 (1:150; Santa Cruz Biotechnology; sc-50715), Collagen I (1:100; Abcam; ab34710), IL-6 (1:50; Novus; NB600-1131), TGF-β1 (1:100; Abcam; ab92486), IL-1β (1:100; Novus; NB600-633), p19 (1:100; Novus; NB200-106), p21 (1:100, Novus NB1001941), Collagen IX (1:500; Abcam; ab134568), COMP (1:200; Abcam; ab231977), Collagen II (1:400; Fitzgerald; 70R-CR008). Tissues sections were washed and incubated with the appropriate Alexa Fluor®-594 conjugated secondary antibody (1:700; Jackson ImmunoResearch Laboratories, Inc., West Grove, PA, USA) for one hour in the dark at room temperature. Sections were washed again with PBS-T (0.4% Triton X-100 in PBS) and mounted with ProLong^®^ Gold Antifade Mountant with DAPI (Thermo Fisher Scientific; P36934). Mounted sections were visualized with Axio Imager 2 (Carl Zeiss Microscopy), using a 10×/0.3 EC Plan-Neofluar (Carl Zeiss Microscopy) objective, X-Cite® 120Q Excitation Light Source (Excelitas Technologies), AxioCam MRm camera (Carl Zeiss Microscopy), and Zen2TM software (Carl Zeiss Microscopy). Exposure settings remained constant across genotypes for each antibody (n=4-6 discs/mouse, 6-7 mice/genotype).

### Digital Image Analysis

All immunohistochemical quantification was conducted in greyscale using the Fiji package of ImageJ (41). Images were thresholded to create binary images, and NP and AF compartments were manually defined using the Freehand Tool. These defined regions of interest were then analyzed either using the Analyze Particles (TUNEL and cell number quantification) function or the Area Fraction measurement.

### Fourier-transform Infrared (FTIR) Spectroscopy

5-μm deparaffinized sections of decalcified caudal disc tissues were collected from STING^+/+^ STING^-/-^ mice (n=1 disc/mouse, 4 mice/genotype) and used to acquire infrared (IR) spectral imaging data using methods previously described (10). Briefly, spectra were collected across the mid-IR region of three consecutive sections/disc. Using the ISys Chemical Imaging Analysis software (v. 5.0.0.14) mean second-derivative absorbances in the amide I (1660 cm-1), collagen side chain vibration (1338 cm-1), and proteoglycan sugar ring (1064 cm-1) regions were quantified and compared in the NP, AF, EP, and vertebrae (VB) of STING^+/+^ STING^-/-^ mice. Significant differences in parameters were assessed by t-test or Mann-Whitney test, where relevant, p < 0.05 was considered significant.

### Spectral Clustering Analysis

Collected spectra were analyzed using K-means clustering analysis in the Eigenvector Solo+MIA software (v. 8.8) to agnostically delineate anatomical regions within the disc, using methods previously reported (10,42). Briefly, during this analysis, regions of IR images are separated into two or more classes, or “clusters”, according to spectral similarity. The K-means partitional clustering method starts with the manually defined selection of K objects that are to be used as cluster targets, where K is determined a priori. During each iteration, the remaining objects (pixels of the spectral image) are assigned to one of these clusters based on the distance from each of the K targets. New cluster targets are then calculated as the means of the objects in each cluster, and the procedure is repeated until no objects are reassigned after the updated mean calculations.

### RNA Isolation, Microarray Assay, and Bioinformatic Analysis

NP and AF tissues were separately micro-dissected from WT and N153S animals under a stereo microscope (Zeiss, Stemi 503) and immediately placed in RNAlater® Reagent (Invitrogen, Carlsbad, CA) as previously described(30). For each animal, tissues from L1-S1 and Ca1-Ca15 were pooled to serve as a single sample (n = 20 discs/animal, 5 animals/genotype). NP and AF tissues were homogenized with a Pellet Pestle Motor (Sigma Aldrich, Z359971), and RNA was extracted from the lysates using an RNeasy® Mini kit (Qiagen). Purified DNA-free RNA was quantified, and quality was assessed using a Nanodrop ND 100 spectrophotometer (Thermo Fisher Scientific) and Agilent 2200 TapeStation (Agilent Technologies, Palo Alto, CA, USA) respectively. The GeneChip Pico kit (Thermo Fisher Scientific) was used to synthesize fragmented biotin-labeled cDNA. Mouse Clariom S gene chips were hybridized with fragmented, biotin-labeled cDNA in 100 μL of hybridization cocktail. Arrays were washed and stained with GeneChip hybridization wash and stain kit using GeneChip Fluidics Station 450 (Thermo Fisher Scientific) and subsequently scanned on an Affymetrix GeneChip Scanner 3000 7G, using Command Console Software (Affymetrix, Santa Clara, CA, USA). CHP files were generated by sst-rma normalization from CEL files and quality control of the experiment was performed in the Transcriptome Analysis Console (TAC) v4.0.2 (Affymetrix). The experimental group was compared to the control group in the TAC, including all probe sets where at least 50% of the samples had a DABG (detected above background) p < 0.05. Inclusion cutoffs were defined at a 2.0-fold change and p-value ≤ 0.05. Significantly differentially up- and downregulated genes from the NP and AF compartments were analyzed using the Overrepresentation Test in PANTHER (43,44).

### Human Microarray Data Analysis

Human sample details are as reported by Kazian et al., and data were obtained from GSE70362 in the GEO database(45). Hierarchical clustering comparing DEGs (inclusion cutoffs of p < 0.05 and 2.0-fold change) in degenerative and non-degenerative samples was conducted with a Euclidean distance < 0.05 cut-off. Each degenerated cluster defined was compared to the aggregated non-degenerated cluster, and the up- and downregulated DEGs identified by each comparison were analyzed for GO biological process enrichment using the PANTHER overrepresentation test, FDR ≤ 0.05.

### Statistical Analyses

Statistical analysis was performed using Prism 9 (GraphPad, La Jolla, CA, USA) with data presented as mean ± standard deviation (SD), p<0.05. For in vivo analyses, data distribution was checked with the Shapiro-Wilk normality test; a Student’s t-test was applied to normally distributed data, and a Mann-Whitney test was applied to non-normally distributed data. Distribution data were compared using a *χ*^2^ test.

## Results

### N153S mice evidence elevated levels of circulating proinflammatory cytokines

To assess the systemic impact of constitutive STING activity, we measured the levels of 19 cytokines and proinflammatory molecules from the plasma of 6-month-old N153S and WT mice using an MSD multiplex assay. N153S mice carry a Sting^1em1Jmin^ mutation resulting in the production of constitutively active STING from the endogenous locus without altering the expression level (29). N153S mice showed significant increases in IL-1β (p = 0.0174) (Fig. 1A), IL-6 (p = 0.0079) (Fig. 1B), TNF-α (p = 0.0360) (Fig. 1C), IFN-γ (p = 0.0074) (Fig. 1D), MIP-1α/CCL3 (p = 0.0025) (Fig. 1E), IL-2 (p = 0.0152) (Fig. 1F), and IL-27/p28/IL-30 (p = 0.0016) (Fig. 1G), MCP-1 (p = 0.0037) (Fig. 1H), and IP-10 (p = 0.0142) (Fig. 1I). We did not observe significant changes in plasma levels of IL-33 (Fig. 1J), IL-5 (Fig. 1K), IL-10 (Fig. 1L), IL-15 (Fig. 1M), MIP-2 (Fig. 1N), or KC/GRO (Fig. 1O); whereas IL-4, IL-9, IL-12/p70, and IL-17p28/IL-30 levels were outside of the assay’s detection limits. These results suggested systemic hypercytokinemia associated with cGAS-STING activation.

**Figure 1.**
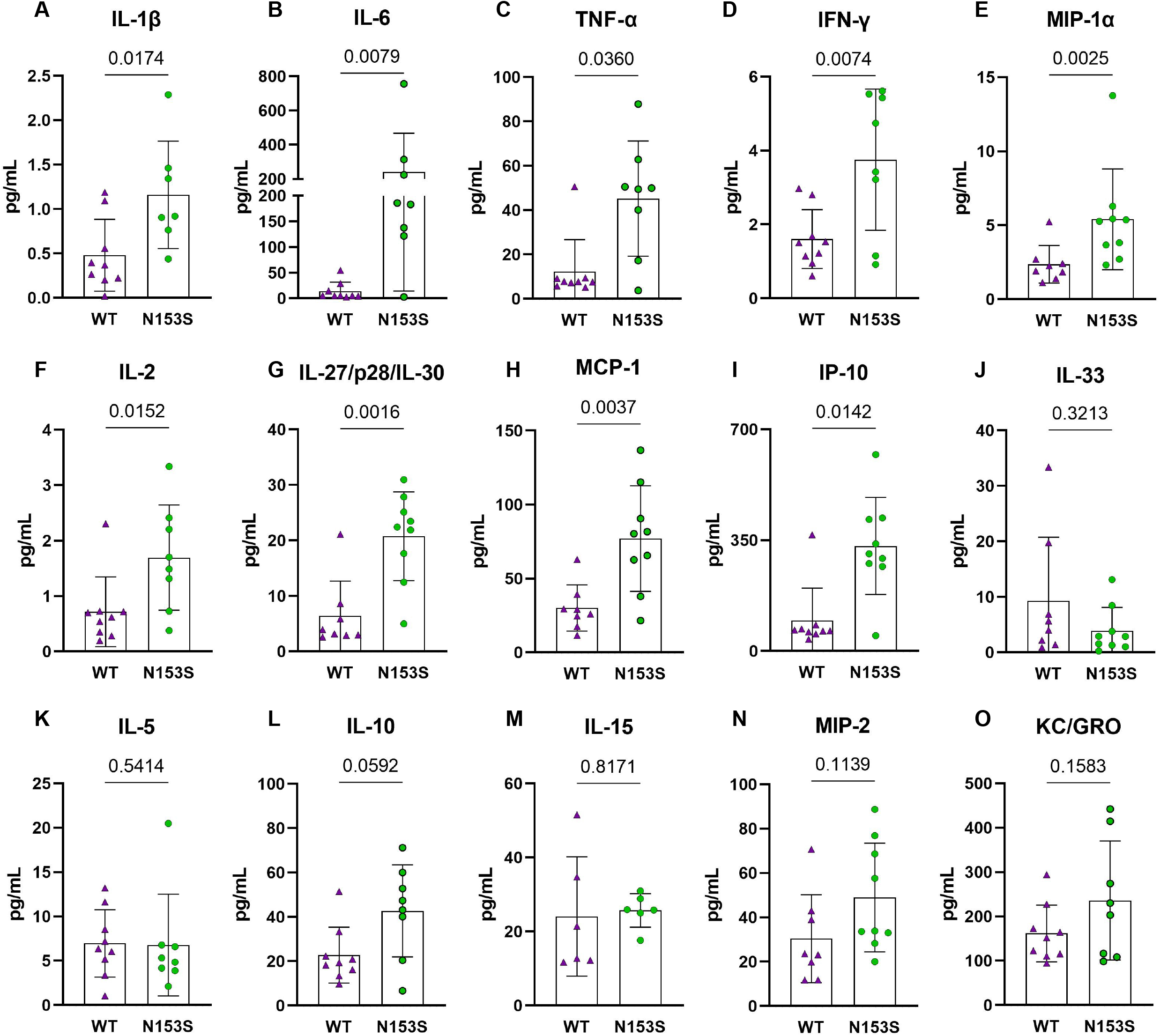
Circulating cytokine levels indicate a proinflammatory environment in N153S mice. Multiplex assay analysis shows a significant increase in the concentrations of (A) IL-1β, (B) IL-6, (C) TNF-α, (D) IFN-γ, (E) MIP-1α, (F) IL-2, (G) IL-27/p28/IL-30, (H) MCP-1, and (I) IP-10 in the plasma of N153S mice. Levels of (J) IL-33, (K) IL-5, (L) IL-10, (M) IL-15, (N) MIP-2, and (O) KC/GRO did not differ between genotypes. Data are shown as mean ± SD. (n = 9 animals/genotype) Significance was determined using an unpaired t-test or Mann-Whitney test, as appropriate.

### Constitutive STING activity compromises trabecular and cortical vertebral bone properties but not knee joints

In models of systemic inflammation induced by DNase II and Ifnar^-^ inactivation, activation of STING pathway has been implicated in pathological bone formation (26). STING deficiency in these mice resolved trabecular bone accrual in long bones and spleens, and transcriptomic analysis indicated STING modulates osteoblast activity, bone matrix remodeling, and osteoclast activity (26). While this offers insight into how STING activity may contribute to the regulation of musculoskeletal tissues, its impact on the spinal column remains to be assessed. To understand the impact of constitutive STING activity on the vertebrae, trabecular and cortical bone morphology were analyzed in the lumbar and caudal spines of WT and N153S mice using micro-computed tomography (μCT). The distal femur was also analyzed for its potential impact on articular joint health. Three-dimensional reconstructions of lumbar (Fig. 2A, A’) and caudal (Fig. B, B’) vertebral motion segments show reductions in the caudal vertebral length (Fig. 2C), lumbar and caudal disc height (Fig. 2D), and the lumbar disc height index (DHI) (Fig. 2E). In the trabecular bone, reductions to the bone volume fraction (BV/TV) (Fig. 2F) and trabecular thickness (Fig. 2G) were observed in N153S caudal but not lumbar vertebrae, while trabecular number (Tb. N.) (Fig. 2H), trabecular spacing (Tb. Sp.) (Fig. 2I), and structure model index (SMI) (Fig. 2J) did not vary across genotypes or region of the spine. Similarly, the bone volume (BV) (Fig. 2K), perimeter (B.Pm.) (Fig. 2L), and area (B. Ar.) (Fig. 2M) were lower in the cortical shell of caudal but not lumbar vertebrae of N153S mice, and the cross-sectional thickness (Cs. Th.) was not impacted (Fig. 2N). Bone mineral density (BMD) was also assessed in the lumbar vertebrae of WT (Fig. 2O) and N153S (Fig. 2O’) animals through the vertebral body (Fig. 2P) and boney endplates (Fig. 2Q); no differences were observed across genotypes. This was consistent with a lack of differences in TRAP (Fig. 2R, R’) or TNAP (Fig. 2S, S’) activity stains. Moreover, the changes observed in the caudal vertebrae did not translate to altered biomechanics, as indicated by the structural (Suppl. Fig. 1A-D) and corresponding material properties (Suppl. Fig. 1E-H). Though ultimate displacement (Suppl. Fig. 1C) was higher in N153S vertebrae, this change was lost when scaled to geometry, as seen from comparable ultimate strain (Suppl. Fig. 1G). It is important to note that trends toward a lower ultimate load (Suppl. Fig. 1A) and higher ultimate stress (Suppl. Fig. 1E) were due to the smaller size of the N153S vertebrae. Taken together, this analysis suggests it is unlikely N153S vertebrae manifested any functional consequences of STING activation. In the knees of WT and N153S mice (Fig. 2T, T’), μCT analysis showed STING activation did not result in osteophyte formation or meniscal ossification. Additionally, 3D analysis of the joint demonstrated comparable BV/TV (Fig. 2U), Tb. Sp. (Fig. 2V), Tb. Th. (Fig. W), and subchondral bone plate thickness (SCBP Th.) (Fig. 2X) between genotypes. Taken together, these results demonstrate a region-specific impact of STING activation on the vertebrae and indicate STING activity reduces caudal trabecular and cortical bone structural metrics without significantly affecting bone function.

**Figure 2.**
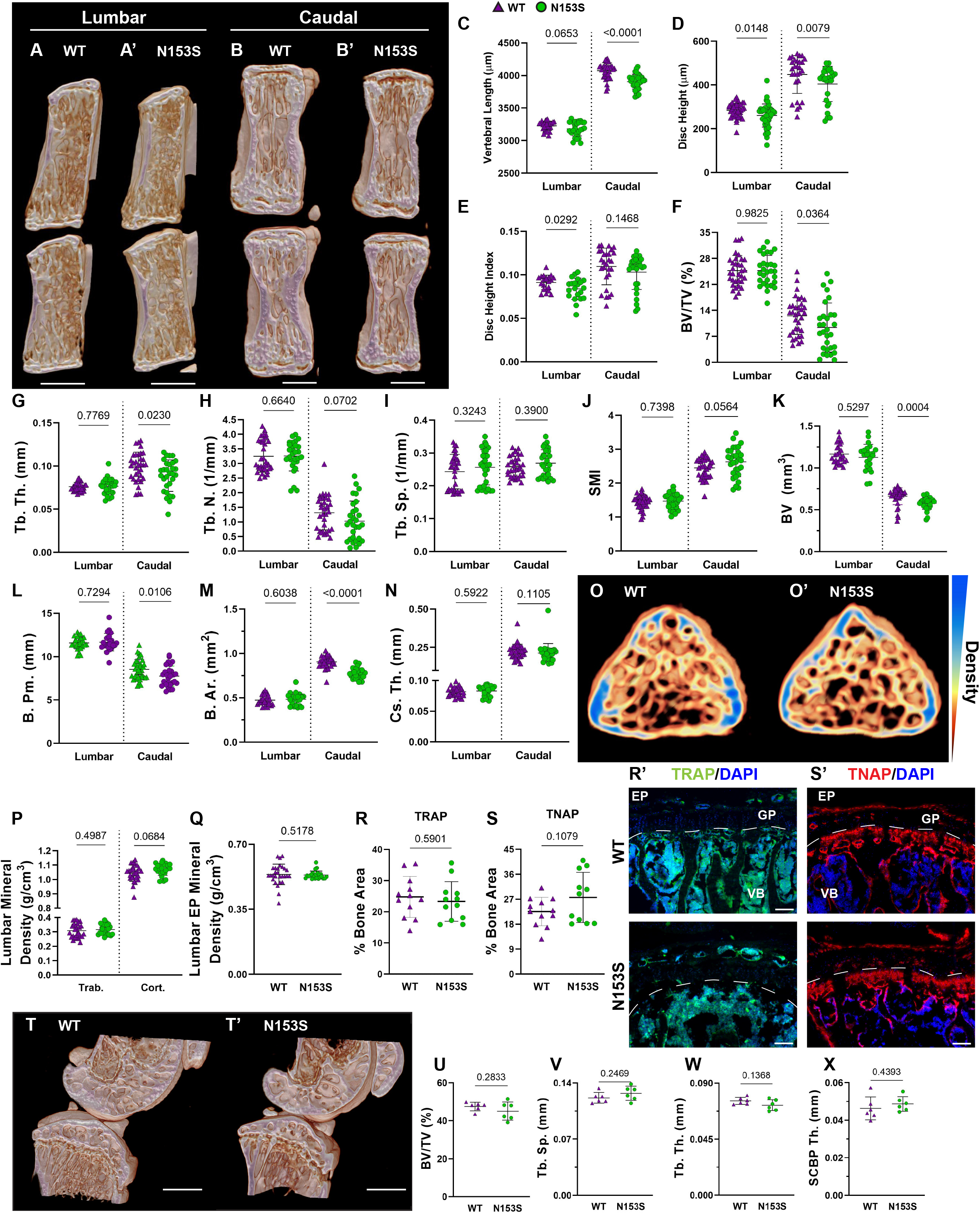
Constitutive STING activity compromises trabecular and cortical vertebral bone properties but not knee joints. (A-B’) Representative μCT reconstructions of the hemi-section of a lumbar (A, A’) and caudal (B, B’) motion segment in a 6-month-old WT and N153S mouse. (C) Vertebral length, (D) disc height, (E) and disc height index (DHI) are shown for lumbar and caudal vertebrae. (F-J) Trabecular bone properties of (F) BV/TV, (G) Tb. Th. (trabecular thickness), (H) Tb. N. (trabecular number), (I) Trab. Sp. (trabecular separation), and (J) SMI (structure model index) are shown for lumbar and caudal vertebrae. (K-N) Cortical bone properties of (K) BV (bone volume), (L) B. Pm. (bone perimeter), (M), B. Ar. (bone area), and (N) Cs. Th. (cross-sectional thickness) are shown. (O, O’) Representative cross-sections of lumbar WT and N153S vertebral bodies, with mineral density indicated on a yellow (less dense) to blue gradient (more dense). (P) Overall trabecular and cortical mineral density and (Q) boney endplate mineral density are shown. Quantitative immunofluorescence of (R, R’) TRAP and (S, S’) TNAP activity in the caudal vertebrae. Dotted lines delineate the growth plate (GP) from the rest of the vertebral body (VB). (T, T’) Representative μCT reconstructions of the medial femoral condyle and tibial plateau. (U) BV/TV, (V) trabecular separation (Tb. Sp.), (W) trabecular thickness (Tb. Th.), and (X) the subchondral bone plate thickness (SCBP Th.) in the hind joints demonstrate no deviations in N153S mice. (n=2 joints/animal, 3 animals/genotype, 6 joints/genotype) Quantitative measurements represent mean ± SD (n=6 lumbar discs and 4 vertebrae/mouse, 7 mice/genotype; n=4 caudal discs and 5 vertebrae/mouse, 7 mice/genotype). (A-B’) Scale bar = 1mm. (O, O’) Scale bar = 250 μm. Significance was determined using an unpaired t-test or Mann-Whitney test, as appropriate.

### Intervertebral discs and articular cartilage of N153S mice do not evidence accelerated degeneration

hSTING-N154S transgenic mice show cartilaginous tissue damage, evidenced by paw swelling, tail shortening, and the loss of ear cartilage (46). Noteworthy, unlike N153S mice, these mice express hN154S, a constitutively active human mutation, through the Rosa locus, resulting in elevated STING activity and higher expression levels of hSTING (46,47). We thereby investigated the histological consequences of constitutive STING activation in the cartilaginous tissues of N153S mice. Safranin O/fast green and hematoxylin staining showed the overall disc tissue architecture was well-preserved in N153S relative to WT mice in lumbar (Fig. 3A, A’) and caudal (Fig. 3B, B’) regions. N153S discs were healthy, as demonstrated by aggrecan-rich NP tissue, vacuolated NP cells, well-organized lamellae in the AF, and a uniform layer of hyaline cartilage with discrete cellular organization in the EP. Modified Thompson scores support these observations, demonstrating no differences in the average (Fig. 3D) NP or AF scores across genotypes in lumbar and caudal regions; however, there were differences in the distribution of lumbar scores and caudal AF scores (Fig. 3C) (37,48–51). These differences did not manifest in any level-by-level changes in the average histological scores (Fig. 3E-F’) and largely do not indicate any trends toward a degenerative phenotype in N153S compared to WT mice. Morphological assessment of the NP health was affirmed at the molecular level, with immunohistological staining showing comparable abundance of aggrecan (Fig. 3G, G’) and the NP-phenotypic marker carbonic anhydrase 3 (Fig. 3H, H’) between N153S and WT mice. Interestingly, despite uncompromised AF morphology, collagen I (Fig. 3I, I’) abundance was lower in the AF of N153S animals. To determine if apoptosis was accelerated by constitutive STING activity in the disc tissues of N153S animals, TUNEL staining was performed (Fig. 3J), demonstrating no increase in apoptosis (Fig. 3J’) and no change in the number of disc cells (Fig. 3J”) in N153S mice. The articular cartilage health in the knee joint was also assessed with toluidine blue (Fig. 3K, K’) and H&E (Fig.3L, L’) staining, with stains showing no damage to the tissue, cell death, or osteophyte formation in N153S mice. When ACS scoring was conducted by blinded graders, N153S and WT animals all received scores of zero, indicating healthy cartilage (38,39). Intact disc architecture, overall retention of core structural proteins, and the lack of evidence supporting increased cell death in N153S discs and knee articular cartilage indicate STING activity does not accelerate disc or articular cartilage degeneration.

**Figure 3.**
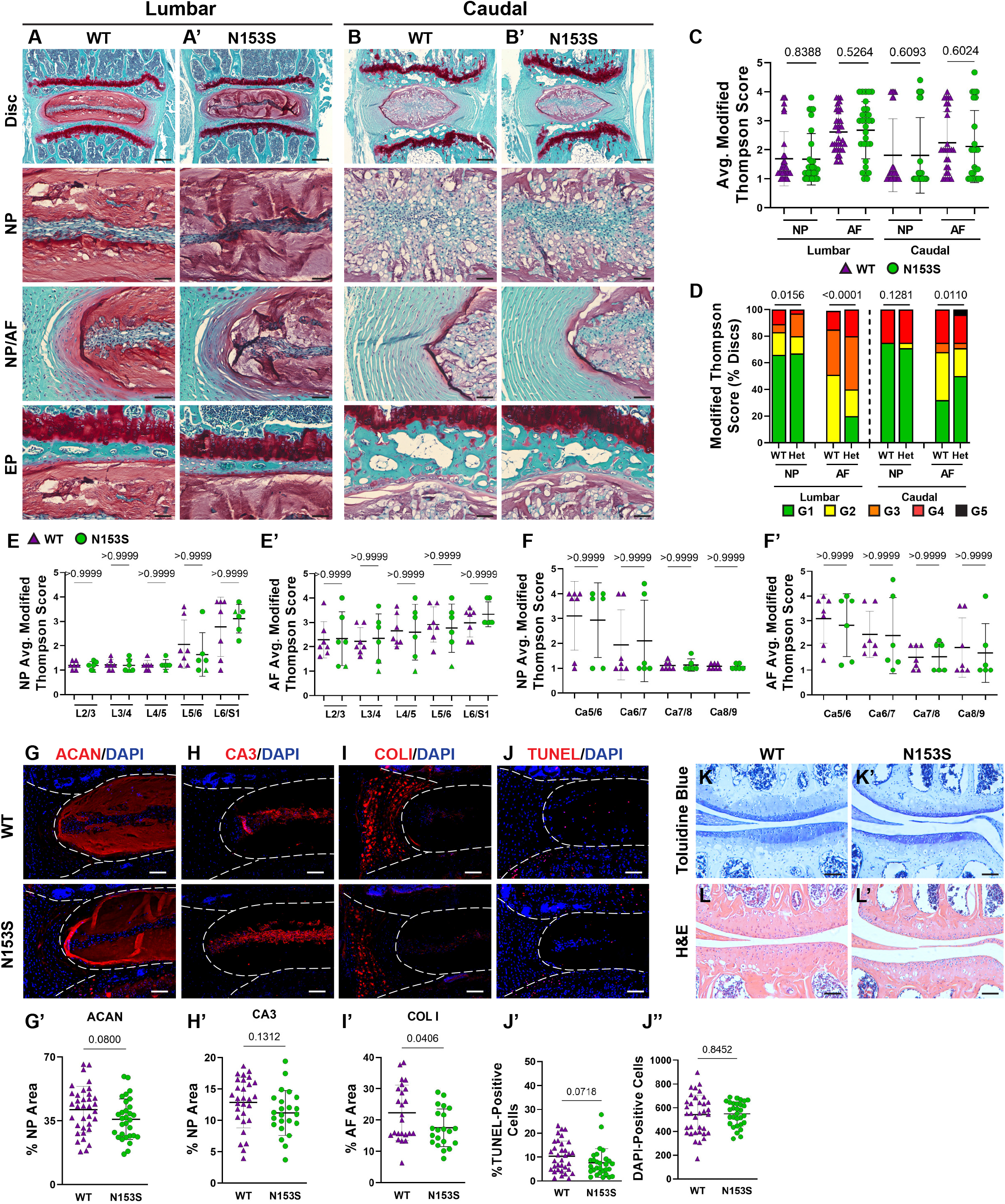
Intervertebral discs and articular cartilage of N153S mice do not evidence accelerated degeneration. (A-B’) 6-month-old WT and N153S mice were analyzed and Safranin O/Fast Green staining of (A, A’) lumbar and (B, B’) caudal discs shows tissue morphology and proteoglycan content (row 1, scale bar= 200 μm) and high magnification images of the NP, EP, and NP/AF tissue boundary (rows 2-4, scale bar= 50 μm) are shown. N153S discs retain vacuolated cells in the NP, demarcation between tissue compartments, and tissue architecture. (C-F’) Histological grading assessment using the modified Thompson scale for (E, E’) lumbar and (F, F’) caudal discs (n=5 lumbar discs/animal, 4 caudal discs/animal, 7 animals/genotype, 35 lumbar and 28 caudal discs/genotype). (C) average and (D) distribution of histological grades in the NP and AF, with higher scores indicating higher levels of degeneration. (E-F’) Level-by-level average grades of NP and AF degeneration. (G-I’) Quantitative immunohistological staining conducted on lumbar discs for: (G) aggrecan (ACAN), (H) carbonic anhydrase 3 (CA3), and (I) collagen I (COL I). (G’-I’) Staining quantification demonstrating no change in ACAN and CA3 abundance and a slight reduction in COL I. (J-J”) TUNEL staining showing apoptotic cells in the NP and AF regions of lumbar intervertebral disc sections from WT and N153S mice. (H, I) Corresponding quantification showing the percentage of TUNEL-positive cells and the number of DAPI-stained nuclei in the entire disc compartment. (K, K’) Toluidine blue staining in the medial femoral condyle and tibial plateau shows no alteration to proteoglycan content and cartilage damage/loss. (L, L’) H&E hindlimb staining showing no apparent cell death or osteophyte formation. (G-L’ scale bar=100 μm) (n=4 discs/animal, 7 animals/genotype, 28 total discs/genotype) (n=1 joint/animal, 3 animals/genotype, 3 joints/genotype) Significance for grading distribution was determined using a χ^2^ test. Significance of average and level-by-level grading and immunohistochemistry was determined using an unpaired t-test or Mann-Whitney test, as appropriate. Quantitative measurements represent mean ± SD.

### SASP induction is not accelerated in the discs of N153S mice

Though the N153S discs did not evidence signs of degeneration, it remained possible for molecular changes to manifest prior to a morphological phenotype or without impacting the disc compartment morphology (19). To assess if this was the case for SASP induction, we conducted immunohistological staining. While. levels of IL-6 were elevated (Fig. 4A, A’) in the NP of N153S mice, compared to WT animals. However, TGF-β (Fig. 4B, B’), IL-1β (Fig. 4C, C’), levels were unchanged across genotypes. Moreover, abundance of p19 (Fig. 4D, D’) and p21 (Fig. 4E, E’) remained unaffected in N153S mice, indicating STING activation does not impact the overall onset of cellular senescence or SASP induction.

**Figure 4.**
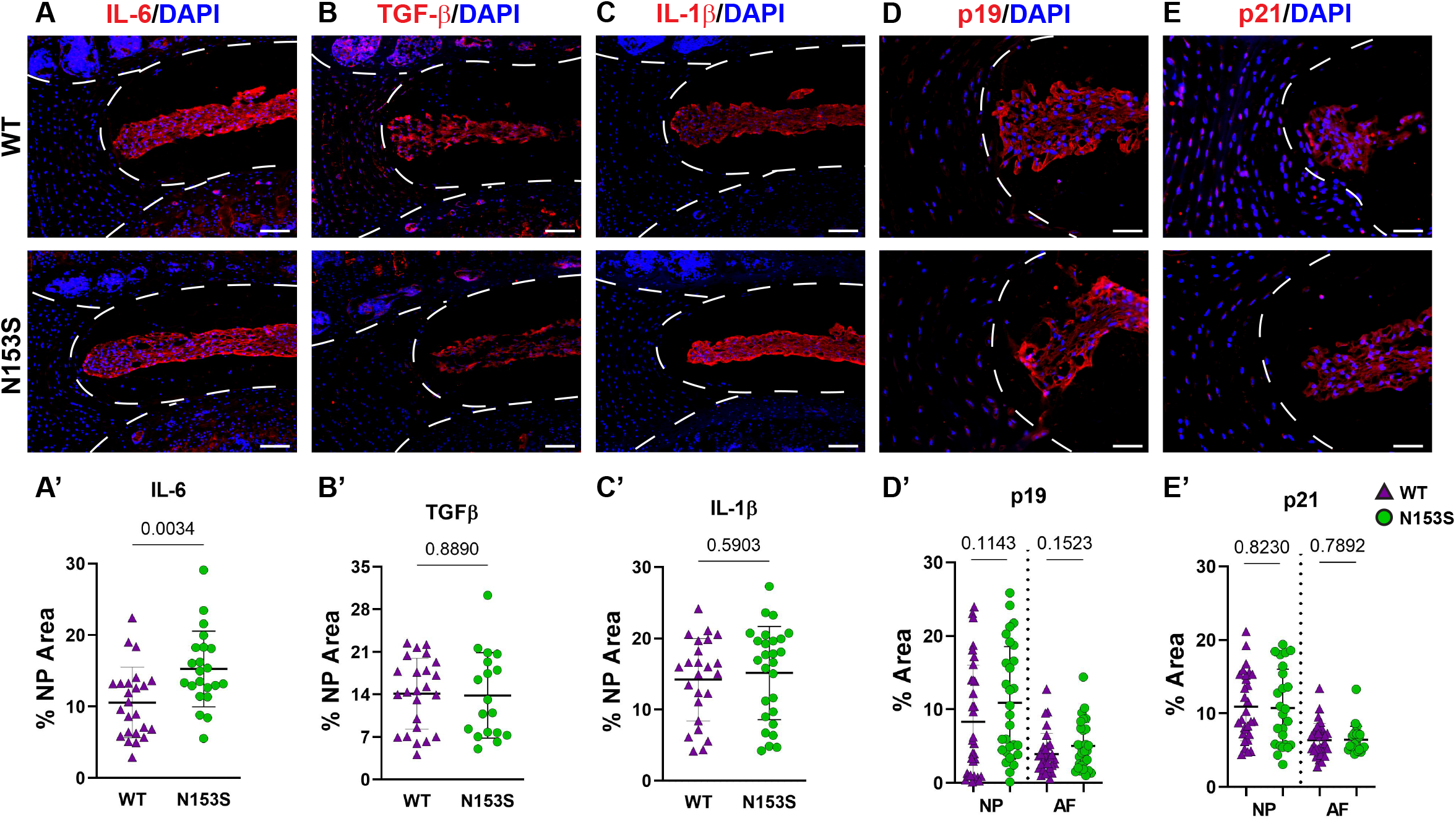
SASP induction is not accelerated in the discs of N153S mice. Quantitative immunohistological staining conducted on 6-month-old WT and N153S lumbar discs for (A, A’) IL-6, (B, B’) TGF-β, (C, C’) IL-1β, (D, D’) p19, and (E, E’) p21. Images were taken at 10x (scale bar=100 μm). (n=4 discs/animal; 7 animals/genotype, 28 total discs/ genotype) Dotted lines demarcate different tissue compartments within the disc. Quantitative data represents the mean ± SD. Significance was determined using unpaired t-test or Mann-Whitney test, as appropriate.

### STING activation results in more prominent transcriptomic changes in the AF than the NP of N153S mice

Microarray analysis was conducted on NP and AF tissues collected from WT and N153S mice to gain insights into transcript-level changes caused by constitutive STING activation. When principal component analysis (PCA) was conducted, WT and N153S NP samples did not cleanly segregate (Fig. 5A); however, hierarchical clustering of differentially expressed genes (DEGs) defined by p ≤ 0.05 and an absolute fold change of two resulted in distinct clustering of WT and N153S NP profiles (Fig. 5B). This profile included 327 DEGs, of which 134 were upregulated and 193 were downregulated, as shown by volcano plot (Fig. 5C). To assess the biological significance of these genes, a statistical overrepresentation test (FDR < 0.05) was conducted in PANTHER, revealing no enrichment for any processes or pathways among DEGs. Among the most up- and downregulated DEGs in the NP were Phkb, Fam13a, and Ric1 and Olfr153, Pard3, and Kpna2, respectively (Fig. 5D). By contrast, AF tissue profiles distinctly clustered by PCA (Fig. 5E) and hierarchical clustering of DEGs (Fig. 5F). Between WT and N153S AF tissues, 1321 DEGs – 856 upregulated and 465 downregulated – were identified (Fig. 5G). As would be expected with STING activation, statistical overrepresentation tests for biological processes in PANTHER revealed immune system processes and their regulation to be those most enriched among upregulated DEGs (Fig. 5H). Some of the select upregulated DEGs included Retnlg, Sll, Lcp1, and Ctsg (Fig. 5H). Downregulated DEGs correlated to the broad ontologies of cellular and metabolic processes (Fig. 5J). When DEGs from these processes were analyzed, the most downregulated genes within the categories were in alignment, including genes such as Esrp2, Mup1, ND6, Adam23, and Serinc4. Taken together, these results indicate that at the transcriptomic level, STING activation more substantially impacts the AF than the NP, with upregulated genes enriching for immune and inflammatory processes.

**Figure 5.**
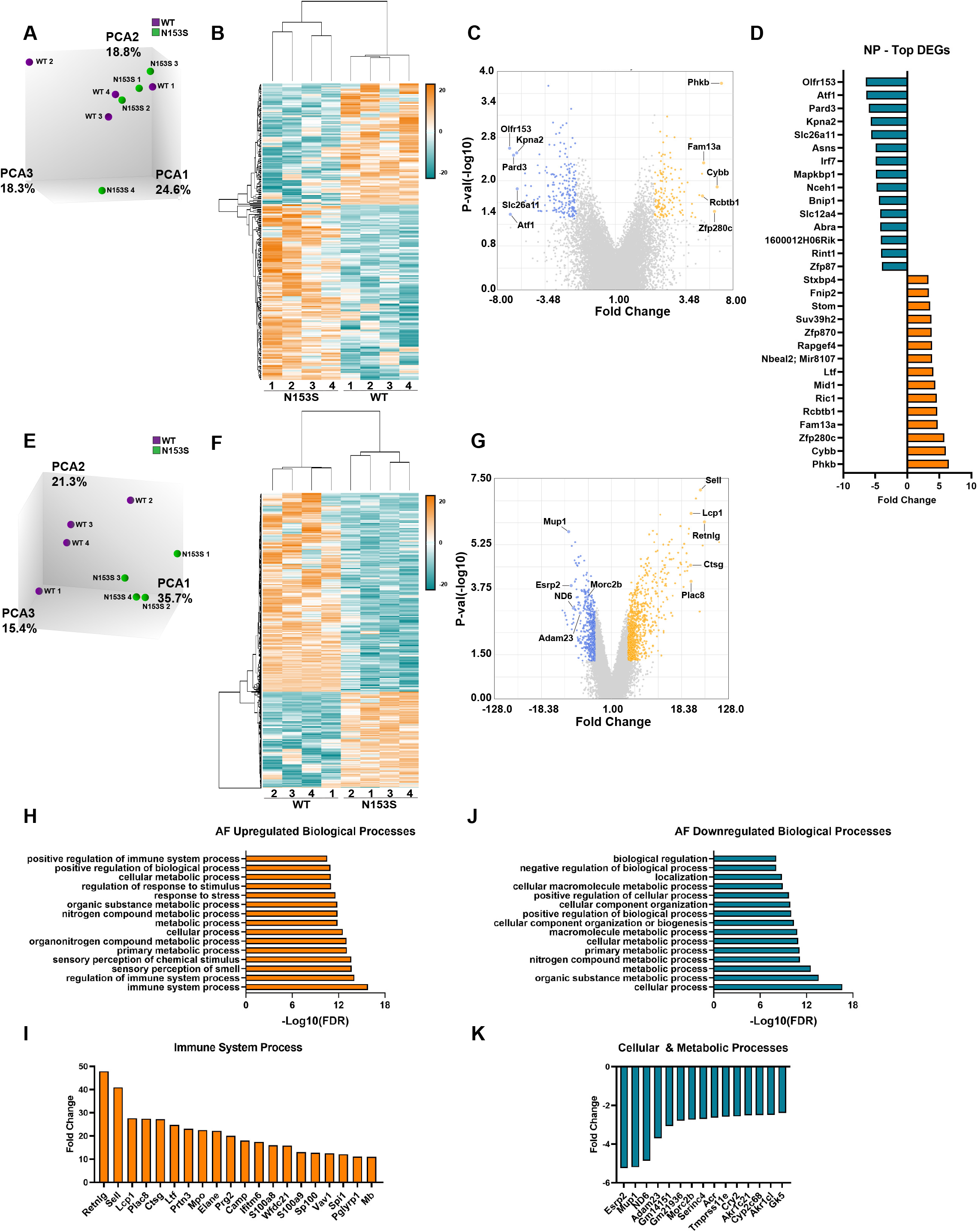
STING activation results in more prominent transcriptomic changes in the AF than the NP of N153S mice. (A) Principal component analysis (PCA) of 6-month-old WT and N153S NP tissues (n = 4 animals/genotype). (B) Heatmap of z-scores for NP DEGs defined by p < 0.05 and 2 < fold change < −2. (C) Volcano plot of DEG p-value plotted against fold change, showing 134 upregulated and 193 downregulated genes in N153S NP tissues. (D) DEGs identified in the NP with the greatest fold change. (E) PCA of 6-month-old WT and N153S AF tissues (n = 4 animals/genotype). (F) Heatmap of z-scores for AF DEGs defined by p < 0.05 and 2 < fold change < −2. (G) Volcano plot of DEG p-value plotted against fold change, showing 856 upregulated and 465 downregulated genes in N153S NP tissues. (H) Representative GO biological processes derived from upregulated genes in N153S AF tissues. (I) DEGs with the greatest fold change within the immune system process GO term. (J) Representative GO biological processes derived from downregulated genes in N153S AF tissues. (K) DEGs with the greatest fold change within the cellular process and metabolic process GO terms.

### Transcriptomic changes in the AF of N153S mice do not recapitulate signatures of human degeneration

To investigate if the gene enrichment signatures observed in the AF tissues of N153S mice captured transcriptomic signatures of human AF degeneration, we analyzed human AF microarray data deposited by Kazezian et al. (GSE70362) (45). Principal component analysis (PCA) demonstrated that human AF samples did not cluster according to Thompson Grade (Fig. 6A). For this reason, we conducted hierarchical clustering of the samples, with a cutoff of Euclidean distance < 0.5 (Fig. 6B) (49). This resulted in four distinct clusters (green boxes) among healthy samples (grades 1 and 2) (n = 7), one mixed cluster (yellow box) consisting of healthy and degenerated samples (grades 1 and 3) (n = 2), and four clusters (pink boxes) among degenerated samples (grades 3, 4, and 5) (n = 15) (Fig. 6B). For subsequent analyses, the healthy clusters were considered as a single group (H), and the mixed cluster (M) and degenerated clusters (D1, D2, D3, D4) were considered as individual clusters (Fig. 6C); this clustering organized accordingly by principal components in three-dimensional space (though this is not captured by the two-dimensional rendering). Clusters D1-D4 and M were compared to cluster H, and the resultant up- and downregulated DEGs (p < 0.05, absolute fold change > 2) from each comparison were analyzed for enrichment of biological processes in PANTHER. Enriched processes were then analyzed against processes enriched in N153S AF tissues. This analysis identified degenerated cluster 2 (D2) as sharing some overlap with the N153S results. Generalized biological and cellular processes were most overrepresented within D2 (Fig. 6D), but processes did include immune and inflammatory response processes which aligned with those identified in the upregulated N153S data (Fig. 6E). However, within these processes, only a few genes were commonly upregulated in human cluster D2 and N153S tissues, including BIRC3, CFD, RSAD2, and ALCAM – none of which are considered inflammatory hallmarks of disc degeneration (Fig. 6F). When downregulated processes were assessed, multiple degenerated clusters showed some overlap with N153S results. This overlap was primarily restricted to generalized cellular, biological, and metabolic processes (Fig. 6G, and there was no overlap in the genes influencing these processes, indicating that in terms of a downregulated transcriptional program, N153S AF tissues do not recapitulate broader immune related processes or signatures of degenerated human AF tissues.

**Figure 6.**
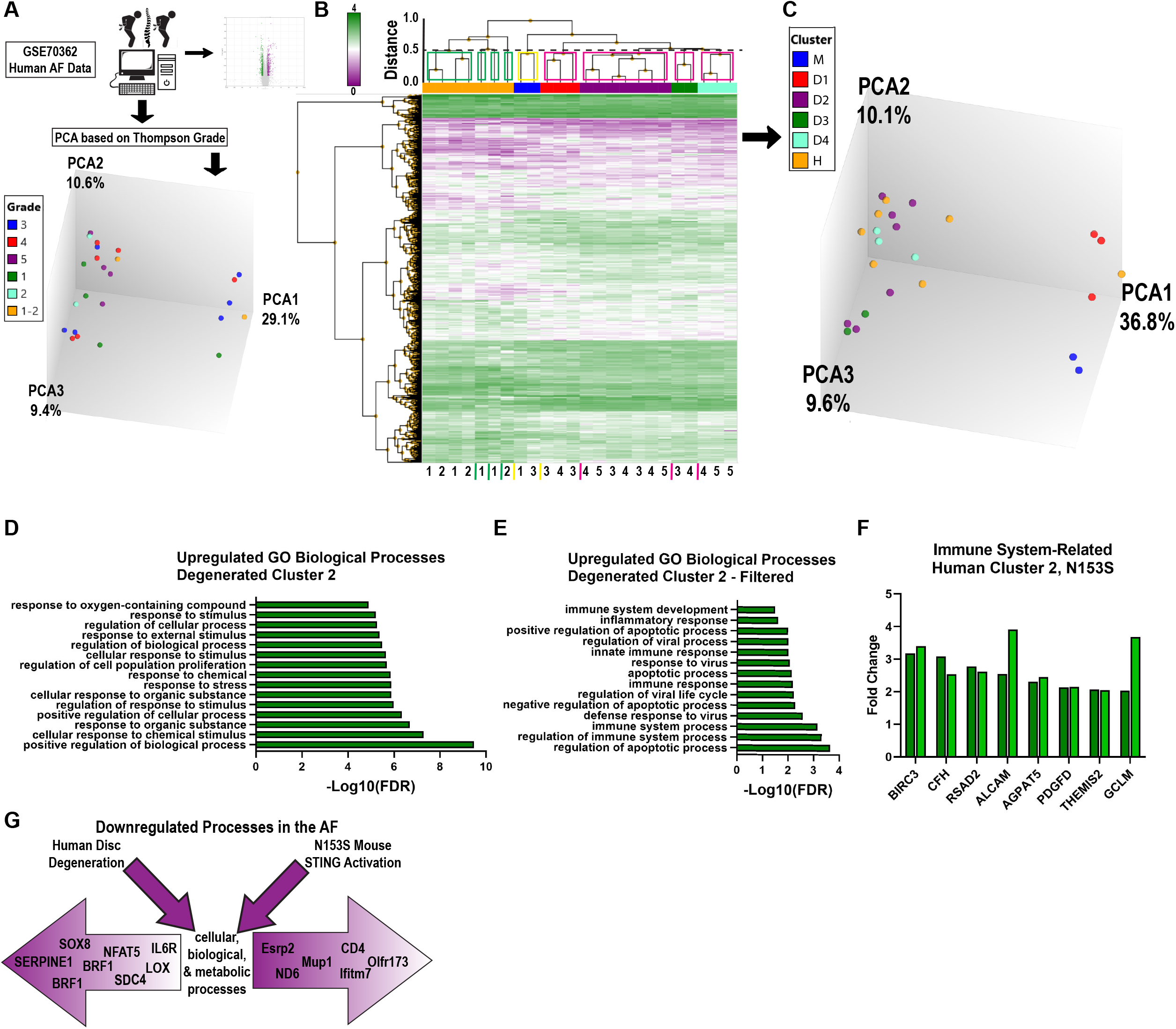
Transcriptomic changes in the AF of N153S mice do not recapitulate signatures of human degeneration. (A) Schematic showing PCA clustering based on Thompson Grades of human AF microarray samples (n=24) deposited in GSE70362. (B) Hierarchical clustering of DEGs, p < 0.05 showed four healthy clusters (green boxes), one ambiguous cluster (yellow box), and four degenerated clusters (pink boxes), with a Euclidean distance cutoff of 0.5. (C) PCA of clusters obtained in B, where all healthy clusters were grouped into a single cluster, H. (D) Representative GO biological processes of upregulated genes between cluster D2 (degenerated) and cluster H (healthy). (E) GO biological processes of upregulated genes between D2 and H that overlap with N153S GO terms related to immune and inflammatory responses. (F) Upregulated DEGs related to an immune response that are common to D2 and N153S transcriptomic profiles. (G) Schematic showing that downregulated genes in the AF of degenerated human AF tissues and N153S AF tissues broadly correlate to cellular, biological, and metabolic processes but diverge in terms of genes contributing to these broad categories of dysregulation.

### Aged STING^-/-^ mice show mild structural changes in the vertebral bone

To investigate the role of the cGAS-STING pathway in an aging context, we analyzed the spinal columns of 16-18-month-old STING^-/-^ mice. Lumbar (Fig. 7A, A’) and caudal (Fig. 7B, B’) vertebrae were assessed using μCT analyses. Caudal vertebrae in STING^-/-^ mice were longer than controls (Fig. 7C), but the length of lumbar vertebrae was comparable between genotypes. In both spinal regions, disc height (Fig. 7D) was unaltered by STING deletion, and the deviations in the caudal vertebral length did not translate to changes in the DHI (Fig. 7E). Three-dimensional analyses of the trabecular bone revealed no changes in the lumbar spine and an increase in the trabecular number (Fig. 7F) in the caudal vertebrae of STING^-/-^ mice. Other trabecular properties of BV/TV (Fig. 7G), trabecular thickness (Fig. 7H), and trabecular separation (Fig. 7I) were consistent between genotypes. Analyses of the cortical shell demonstrated reduced bone volume (Fig. 7J), cross-sectional thickness (Fig. 7K), and bone area (Fig. 7L) in the caudal vertebrae of STING^-/-^ mice and no changes in the lumbar vertebrae. As seen in the N153S model, the consequences of STING deletion in the vertebrae are region-specific, with the caudal vertebrae demonstrating significant alterations in the cortical shell and trabeculae.

**Figure 7.**
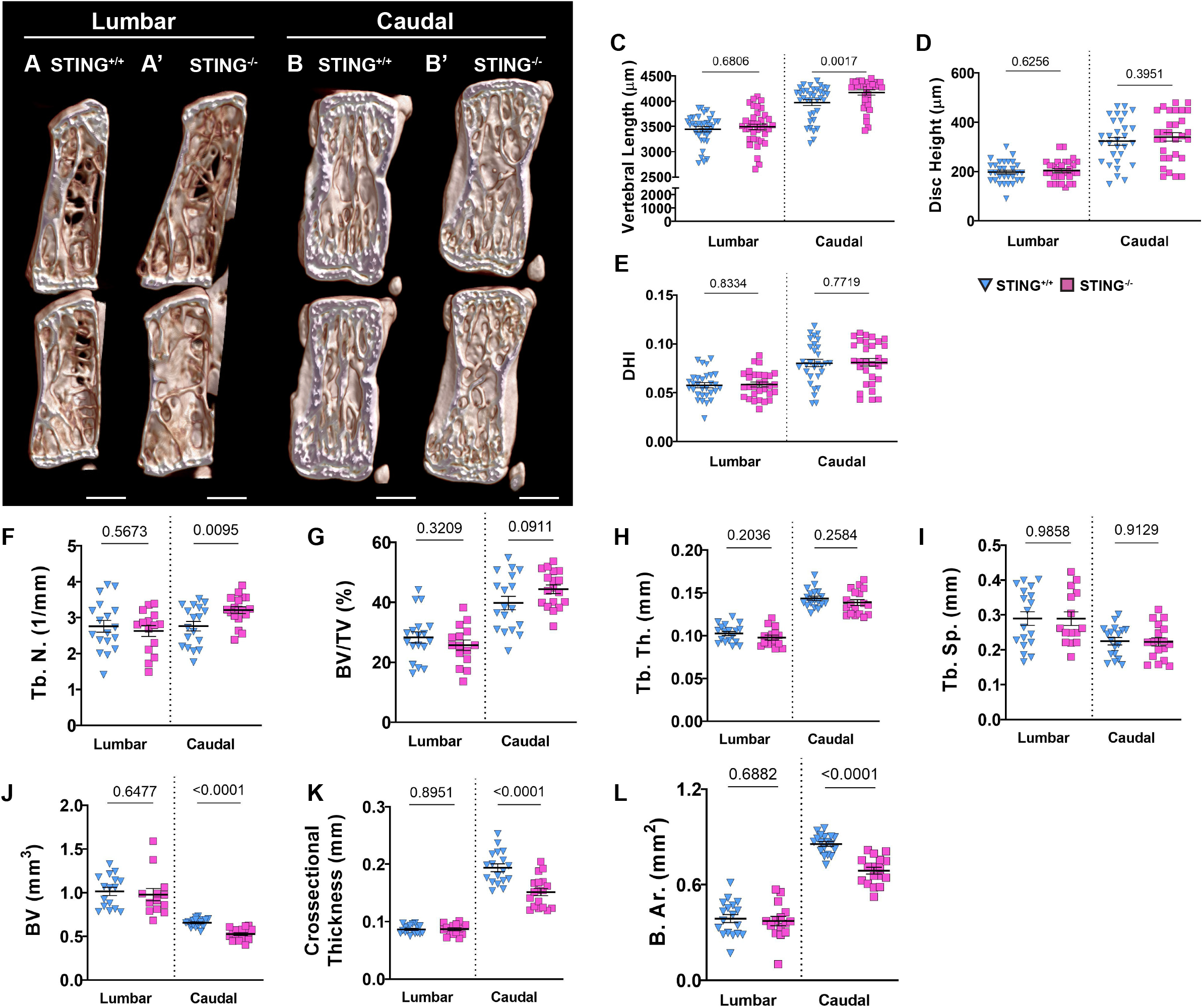
Aged STING−/− mice show mild structural changes in the vertebral bone. (A-B’) Representative μCT reconstructions of the hemi-section of a lumbar (A, A’) and caudal (B, B’) motion segment in a 16-18-month-old STING^+/+^ and STING^-/-^ mouse. (C) Vertebral length, (D) disc height, (E) and disc height index (DHI) are shown for lumbar and caudal vertebrae. (F-I) Trabecular bone properties of (F) Tb. N. (trabecular number), (G) BV/TV, (H) Tb. Th. (trabecular thickness), and (I) Trab. Sp. (trabecular separation) are shown for lumbar and caudal vertebrae. (J-L) Cortical bone properties of (J) BV (bone volume), (K) Cs. Th. (cross-sectional thickness), and (K) B. Ar. (bone area) are shown. Quantitative measurements represent mean ± SD (n=6 lumbar discs and 5 vertebrae/mouse, 6 mice/genotype; n=6 caudal discs and 5 vertebrae/mouse, 6 mice/genotype). (A-B’) Scale bar = 1mm. Significance was determined using an unpaired t-test or Mann-Whitney test, as appropriate.

### Aging STING^-/-^ mice do not evidence better morphological attributes and SASP status in disc

We assessed the impact of STING inactivation in delaying the onset of age-related degenerative changes in the disc. Safranin O/fast green and hematoxylin staining on average did not demonstrate better-preserved tissue architecture in the lumbar (Fig. 8A, A’) or caudal (Fig. 8B, B’) discs of 16-18-month-old STING^-/-^ mice relative to STING^+/+^ mice. Quantitative evaluation using the Modified Thompson grading scheme demonstrated slight differences in the distribution (Fig. 8C) of lumbar NP scores, where no STING^-/-^ discs scored a 3 or 4, and in caudal AF scores, for which distribution deviations did not consistently demonstrate improved or worsened disc health, as STING^-/-^discs had higher proportions of grades 1 and 4. Despite these differences in score distributions, cumulative average grades (Fig. 8D) did not vary across genotypes for either spinal region, and level-by-level analyses (Fig. 8E-F’) did not indicate level-specific deviations, with the exception of the NP at L6/S1, where STING^-/-^ discs averaged lower scores than controls.

**Figure 8.**
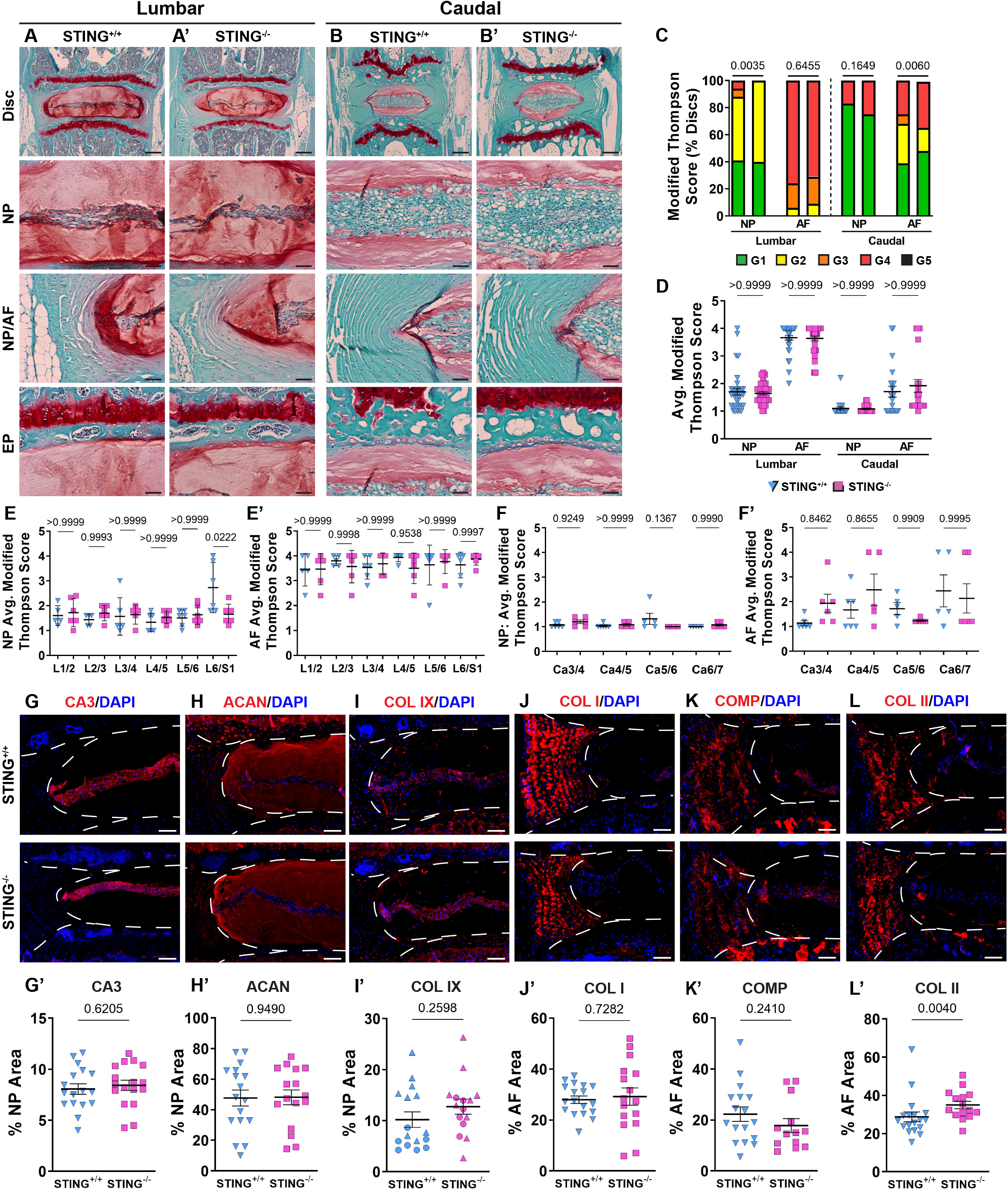
STING^-/-^ discs do not evidence an improved morphological or molecular phenotype. (A-B’) 16-18-month-old STING^+/+^ and STING^-/-^ mice were analyzed and Safranin O/Fast Green staining of (A, A’) lumbar and (B, B’) caudal discs shows tissue morphology and proteoglycan content (row 1, scale bar= 200 μm) and high magnification images of the NP, EP, and NP/AF tissue boundary (rows 2-4, scale bar= 50 μm) are shown. (C-F’) Histological grading assessment using the modified Thompson scale for (E, E’) lumbar and (F, F’) caudal discs (n=5 lumbar discs/animal, 4 caudal discs/animal, 6 animals/genotype, 30 lumbar and 24 caudal discs/genotype). (C) Distribution of and (D) average histological grades in the NP and AF, with higher scores indicating higher levels of degeneration. (E-F’) Level-by-level average grades of NP and AF degeneration. (G-L’) Quantitative immunohistological staining conducted on lumbar discs for (A, A’) CA3, (B, B’) ACAN, (C, C’) collagen IX (COL IX), (J, J’) COL I, (K, K’) collagen oligomeric matrix protein (COMP), and (L, L’) collagen II (COL II). (n=2-4 discs/animal; 6 animals/genotype, 12-24 total discs/genotype) Dotted lines demarcate different tissue compartments within the disc. Significance for grading distribution was determined using a χ^2^ test. Significance of average and level-by-level scores and quantified immunohistochemistry was determined using an unpaired t-test or Mann-Whitney test, as appropriate. Quantitative measurements represent mean ± SD.

At the molecular level, immunohistochemical staining showed carbonic anhydrase 3 (Fig. 8G, G’), aggrecan (Fig. 8H, H’), and collagen IX (Fig. 8I, I’) to be similarly abundant in the NP tissues of STING^-/-^ and STING^+/+^ mice. Abundance of key structural proteins collagen I (Fig. 8J, J’) and cartilage oligomeric matrix protein (COMP) (Fig. 8K, K’) was unchanged in the AF of STING^-/-^ discs; however, these discs showed increased abundance of collagen II (Fig. 8L, L’).

To further assess the overall collagen architecture and content of these discs, Picrosirius Red-stained discs from the lumbar (Suppl. Fig. 2A, A’) and caudal (Suppl. Fig. 2B, B’) spine were visualized under polarized light. Across both genotypes, the AF contained highly organized collagen fibrils, and no staining was detected in the NP. Quantitative analysis showed no differences in collagen fiber thickness distribution (Suppl. Fig. 2C), indicating comparable rates of collagen turnover in knockout and wildtype discs. Fourier transform infrared (FTIR) spectroscopic imaging was used to assess the chemical composition of STING^-/-^ and STING^+/+^ discs. Spectral clustering that groups and classifies individual pixels based on their chemical compositions showed no differences in the chemical composition of knockout and wildtype motion segments (Suppl. Fig. 2D, D’). Average second derivative spectra were generated and analyzed for each disc compartment an the vertebrae at representative peaks for chondroitin sulfate (1064 cm^-1^), cell-associated proteoglycans (1156 cm^-1^), collagen (1338 cm^-1^), and total protein (1660 cm^-1^)(Supp. Fig. 2E, E’). Corresponding chemical maps (Suppl. Fig. 2F-I’), quantified for each molecular class in each tissue (Suppl. Fig. 2F”-G”), revealed only a slight reduction in the collagen content of the NP, without any significant changes to chondroitin sulfate, cell-associated proteoglycans, or total protein content. Taken in context with the histological evaluation of STING^-/-^ and STING^+/+^ mice, this chemical analysis indicates STING inactivation does not confer an advantage to any of the disc compartments during aging.

Again considering the possibility that a molecular phenotype may manifest in the absence in a morphological phenotype, we conducted immunohistological staining for known SASP markers to determine if the loss of STING attenuated delayed the onset of senescence. Interestingly, IL-1β was more abundant in knockout discs than in wildtype discs (Suppl. Fig. 3A, A’), but IL-6 (Suppl. Fig. 3B, B’), TGF-β (Suppl. Fig. 3C, C’), p19 (Suppl. Fig. 3D, D’) and p21 (Suppl. Fig. 3E, E’) were unchanged, indicating loss of STING does not delay cell senescence or SASP onset. Taken together, our analyses of N153S and STING^-/-^ mice for the first time demonstrate that the cGAS-STING pathway is not a critical regulator of senescence and consequent degeneration of the intervertebral disc in vivo.

## Discussion

Cellular senescence is well-established as a feature of aging and degeneration in the human intervertebral disc as well as various mouse models (13,17,19,52). Of interest, cellular senescence is shown to destabilize the nuclear membrane, resulting in the release of cytosolic chromatin fragments and consequent activation of the cGAS-STING pathway (24). This DNA-sensing pathway promotes SASP and mediates type-I interferon inflammatory responses to foreign viral and bacterial DNA as well as self-DNA(22). Cursory studies of NP cells and in vivo injury models have implicated the cGAS-STING pathway as an active inducer of senescence in the disc; however, this pathway in the disc is yet to be explored independent of chemical or mechanical insult (27,28,53–55). In this study, we clearly show STING is not a critical mediator of senescence onset and consequent degeneration in the disc. Detailed analyses of the spinal columns of 6-month-old heterozygous N153S mice with a gain-of-function mutation in the Sting1 gene and 16-18-month-old STING^-/-^ mice revealed healthy disc tissues in both genotypes with some compromised features of the vertebrae.

The gain-of-function STING mutation resulted in a significant increase in the plasma concentration of pro-inflammatory molecules IL-1β, IL-6, TNF-α, IFN-γ, MIP-1α/CCL3, IL-2, IL-27/p28/IL-30, MCP-1, and IP-10 in N153S mice. N153S animals have also been shown to overexpress IL-1β, IL-6, MIP-1α in their lungs as an attribute of the vasculopathy they experience (31). The increased systemic cytokine concentrations are indicative of the hypercytokinemia that occurs as a consequence of cGAS-STING activity; as an immune and antitumor response, this cytokine “storm” is capable of causing cell toxicity and death, which is advantageous as a defense mechanism but otherwise detrimental (22,56). These results underscored an environment of systemic inflammation, which could have broader consequences for the health of spinal column. One such consequence may be the reduction in bone volume observed in the trabecular and cortical bone of the caudal vertebrae in N153S mice. Although previous studies correlated STING expression with elevated osteoblastic activity in a model of DNA damage-induced inflammation (DNaseII, IFNR^-/-^), IL-1β, IL-6, TNF-α stimulate osteoclast differentiation, which may explain the compromised bone quality in N153S mice (26,57,58). Of note, recent work demonstrated STING induction through DMXAA treatment in mice attenuates cancer-induced osteoclast differentiation in the femur through type I interferon signaling and that STING induction inhibited bone fracture-induced pain by suppressing nociceptor excitability (59). These results show an opposite effect of STING activation on bone cells in the femur than that observed in the vertebrae in the present study; one possible explanation for such is the impact of proinflammatory signaling in the absence of direct insult to the tissue as opposed to in the context of injury or diseased tissue microenvironment.

Further, despite the proinflammatory circulating cytokine profile and mild vertebral phenotype, histological analyses demonstrated that STING activation did not accelerate a degenerative phenotype in the NP, AF, EP, or articular cartilage. This was somewhat surprising considering arthritis is a symptom of SAVI, the disease recapitulated by the N153S mutation (60). The findings in the disc do however align with the recent work of Gorth et al. in hTNF-α overexpression mouse models and contribute to a growing body of evidence demonstrating the marginal impacts of systemic inflammation on overall disc health (30,61). Other in vivo models of systemic inflammation, including high fat diet-induced obesity, have shown disc architecture is not negatively impacted by inflammatory conditions that are not local (62,63). As seen in the structurally intact discs of these models, N153S NP tissues retained healthy, vacuolated cells expressing the NP-phenotypic marker CA3 and an aggrecan-rich matrix. Unlike the hTNF-α models, the AF was structurally uncompromised by constitutive STING activation, demonstrating a reduction in collagen 1 abundance without impact on the organization or structure of the tissue. However, at the transcript level, the N153S AF did demonstrate a more significant response to STING activation, upregulating over 100 genes related to an immune response, including il1b, il1rn, ifitm6, cdk1, ifitm3, and tnfaip2. This gene signature, however, did not tightly correlate to one associated with disc degeneration in human AF tissues. Marginal overlap between the results of N153S and degenerated human AF tissues indicates the possibility for STING activation to eventually facilitate the degenerative process, but at the time of analysis, STING activation did not promote a transcriptional program associated with disc degeneration. Considering the NP tissues of N153S animals, despite an increase in IL-6, the overall onset of senescence and induction of SASP were not accelerated by STING gain-of-function, as indicated by unaltered TGF-β, IL-1β, and p19 levels. Together this analysis supports that the NP is largely isolated from the consequences of systemic inflammation and that systemic inflammation does not necessarily cause disc degeneration.

Although STING gain-of-function did not expedite degenerative structural changes or the induction of senescence in the disc, it remained possible that in an aging context – when senescence is known to occur – the deletion of STING would be protective to the spinal column by delaying the onset of senescence. μCT analyses of 16-18-month-old STING^-/-^ vertebrae showed that the loss of STING in the caudal vertebrae resulted in longer bones with fewer trabeculae and an overall reduction in the structural properties of the cortical shell. This finding is interesting in that cortical thinning and reduced bone mass have been associated with systemic inflammation clinically and in mouse models, but STING^-/-^ tissues have been shown to express lower levels of inflammatory markers than STING^+/+^ tissues (30,64). One possible explanation for this contradiction is the apparent impact of STING on the balance between osteoblast and osteoclast activity, as suggested by Baum et al. (26).

Similarly to the N153S model, histological and spectroscopic analyses of the discs revealed STING loss did not improve aging outcomes. Aged STING^-/-^ and STING^+/+^ discs maintained healthy disc architecture and cellularity, as reflected by low Modified Thompson scores and comparable levels of key structural proteins, including aggrecan, collagen IX, collagen I, and COMP. STING^-/-^ discs did evidence higher abundance of collagen II; however, collagen fiber thickness distribution and chemical mapping demonstrated overall consistency across genotypes. This was also true for SASP markers IL-6, TGF-β, and p19, with STING^-/-^ discs showing increased levels of IL-1β. These results strongly support that STING is not a critical effector of senescence in the natural aging and degeneration context. Recent work has demonstrated cGAS-STING activation in a LPS vertebral injury-induced disc degeneration model; however, this activation was only achieved with the inflammatory stimulus of LPS and by breaching the integrity of the EP, enabling immune cell infiltration into the disc compartment (27). Similarly, a recent study showed the degenerative effects of needle puncture injury to rat discs were reduced when STING activation was attenuated by limiting mitochondrial DNA release from damaged mitochondria resulting from calcium overload (28). It is likely, however, that in the context of this acute herniation injury, infiltrating scavenging immune cells such as neutrophils and macrophages were the primary effectors of the cGAS-STING response and responsible for promoting inflammation; a similar immune response has been documented in the Tg197 mouse model, which overexpresses hTNFα and is predisposed to spontaneous disc herniations (61). Our work clearly demonstrates that in the absence of chemical or traumatic mechanical insult, STING activity neither accelerates nor attenuates the onset of cellular senesce and the degeneration cascade in the intervertebral disc. Accordingly, while the cGAS-STING pathway is a critical modulator of cellular senescence in other systems, it is unlikely to be an effective target for mitigating degeneration and cell senescence in the disc.

## Data Availability

The datasets generated and analyzed for this study can be found in the GEO database (GEO accession number GSE188909).

## Author Contributions

Study design: OKO, MVR. Study conduct: OKO, CK, JAC. Data Analysis: OKO, CK, JAC. Data interpretation: OKO, MVR, JAC. Drafting manuscript: OKO, MVR, JAC. Approving final version of the manuscript: OKO, CK, JAC, MVR.

## Acknowledgments

We thank the Sidney Kimmel Cancer Center Cancer Genomics Facility of Thomas Jefferson University, and Dr. Ryan Tomlinson for help with biomechanical analysis. We thank Drs. Shelley Berger and Zhixun Dou at the University of Pennsylvania for providing tissues from STING^+/+^ and STING^-/-^ mice.

---

**Supplementary Figure 1.**
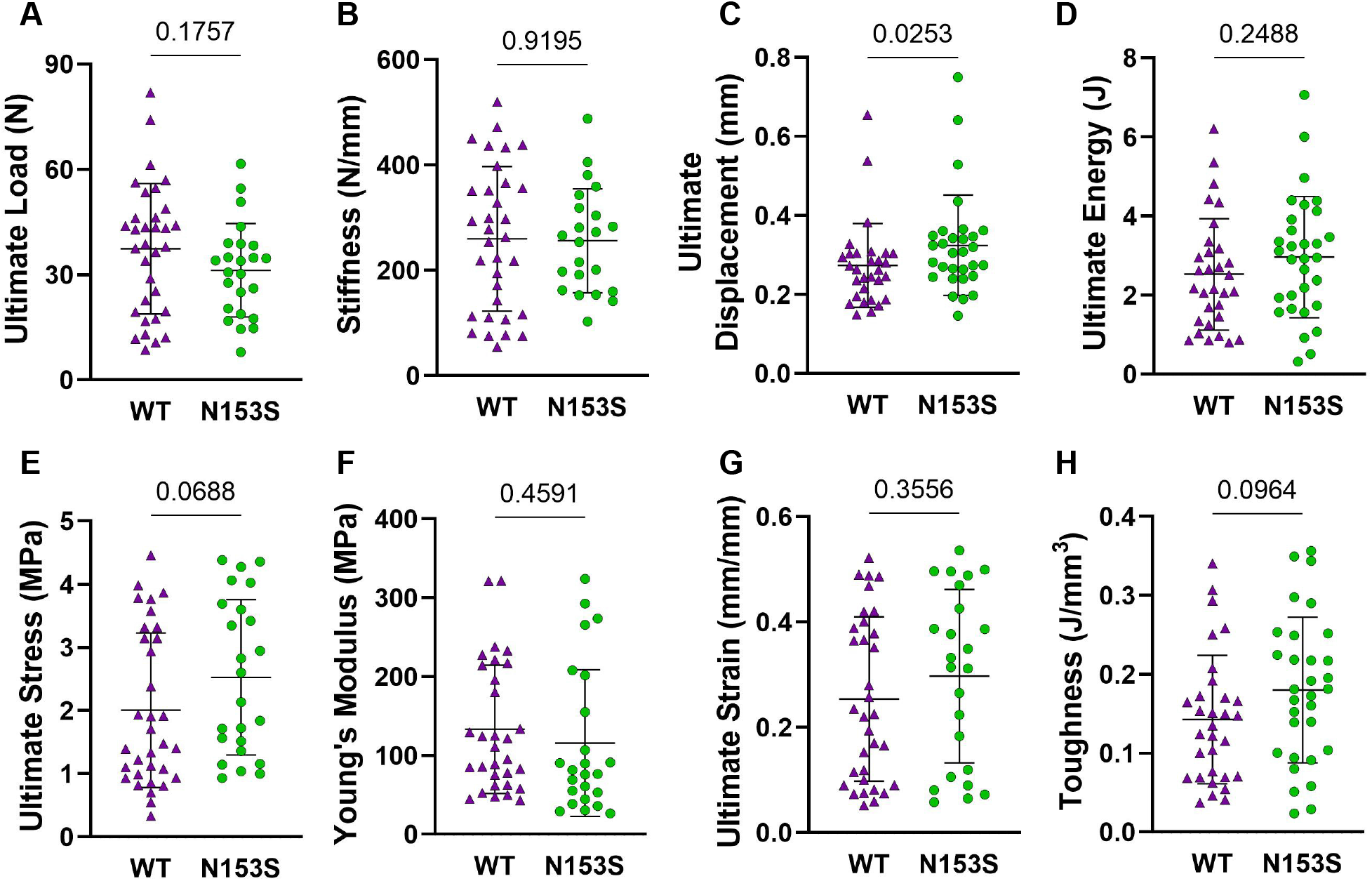
Changes in N153S vertebral architecture do not lead to functional changes. WT and N153S caudal vertebral bone structural properties of (A) ultimate load, (B), stiffness, (c) ultimate displacement, and (D) ultimate energy and their corresponding material properties of (E) ultimate stress, (F) Young’s Modulus, (G) ultimate strain, and (H) toughness, as determined by compression testing. (n=2-3 caudal vertebrae/animal, 12 animals/genotype, 27-32 vertebrae/genotype). Data are represented as the mean ± SD. Significance was determined using an unpaired t-test or Mann-Whitney test, as appropriate.

**Supplementary Figure 2.**
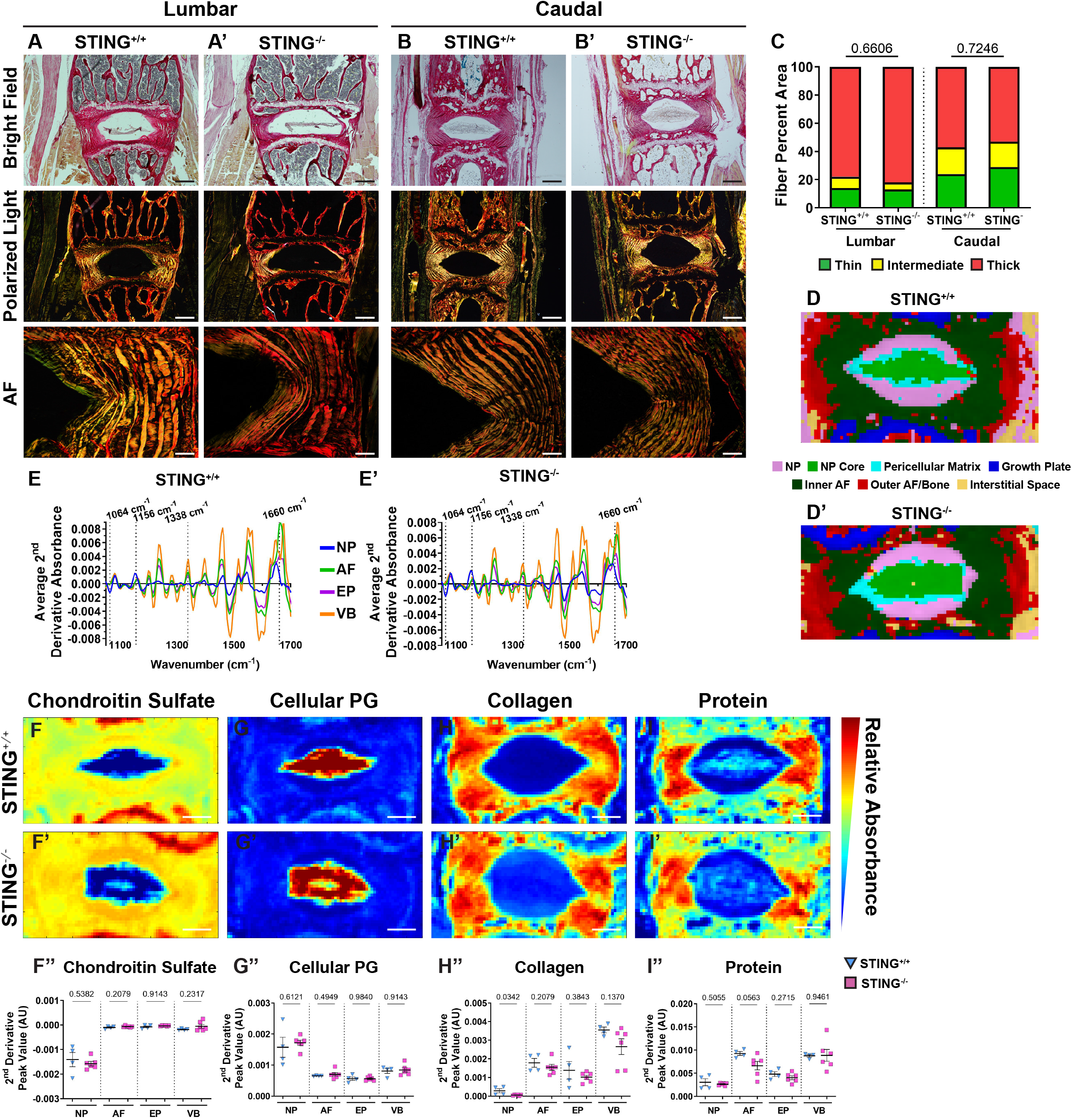
STING deletion does not alter collagen structure or chemical composition in the aging mouse disc. (A-B’) Picrosirius Red staining of 16-18-month-old (A, A’) lumbar and (B, B’) caudal discs showing collagen organization of the AF in the bright field (top row) and collagen fiber distribution under polarized light (middle and bottom rows) (scale bar= 100 μm). (C) Quantification of fiber thickness distribution (n=5 lumbar discs, 4 caudal discs/animal, 6 animals/genotype, 30 lumbar discs/genotype, 24 lumbar discs/genotype). (D, D’) Average second derivative spectra, inverted for positive visualization, of the NP, AF, EP, and vertebrae (VB) of (H, H’) of 16-18-month-old STING^+/+^ and STING^-/-^ mice (n=1 lumbar disc/animal, 4 animals/genotype, 4 total discs/genotype). (E, E’) Spectral cluster analysis images (Scale bar = 200 μm). (F-H’) Chemical maps and (F”-H”) quantification of mean second derivative peaks for (F-F”) chondroitin sulfate (1064 cm^-1^), (G-G”) collagen (1338 cm^-1^), and (H-H”) total protein (1660 cm^-1^) content. Significance between fiber distribution was determined using a χ2 test. AU: arbitrary units. Quantitative measurements represent mean ± SD. Significance of chemical components was determined using an unpaired t-test or Mann-Whitney test, if data were not normally distributed.

**Supplementary Figure 3.**
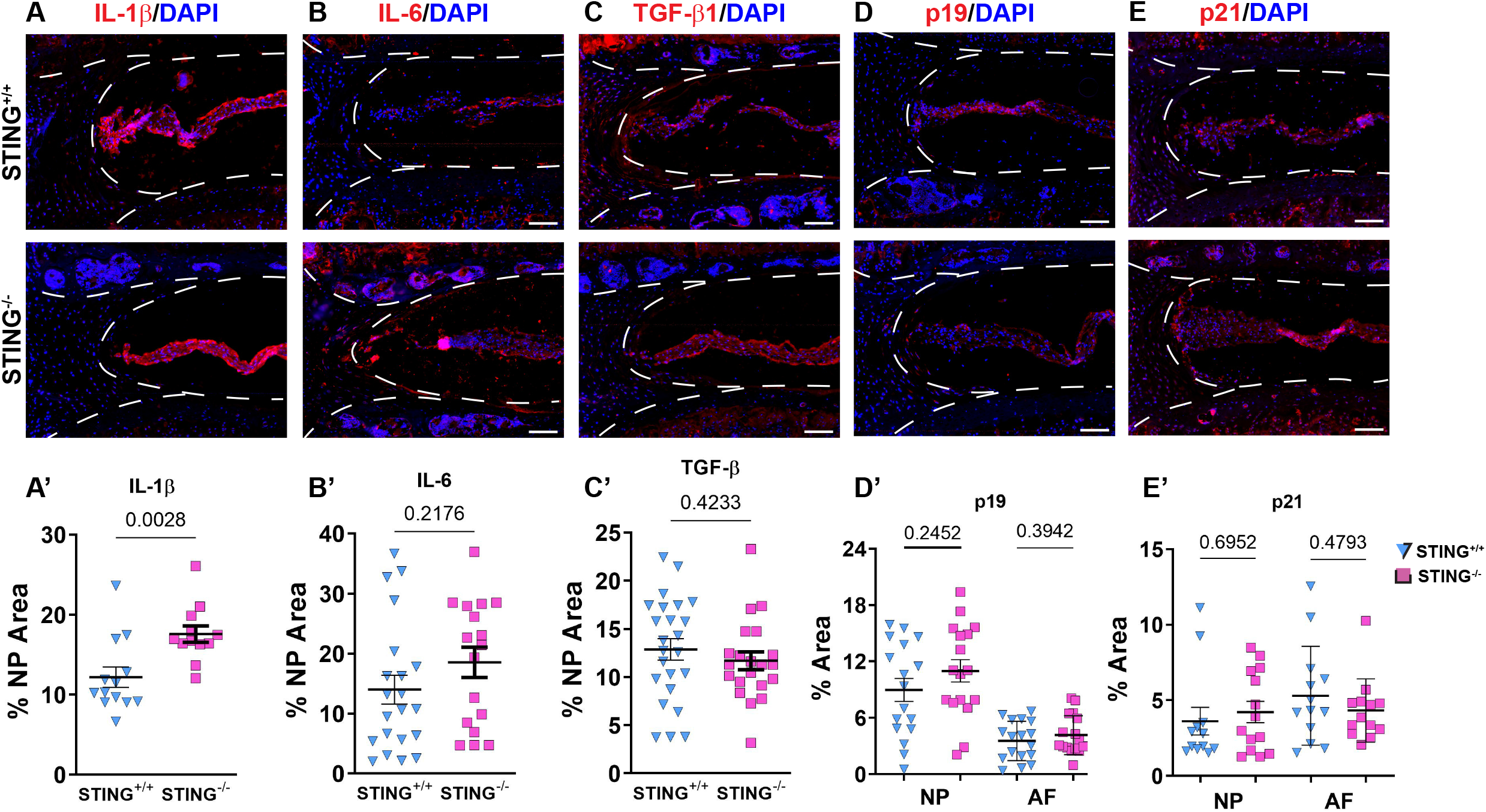
SASP induction is not delayed in STING^-/-^ mice. (A-D’) Quantitative immunohistological staining of 16-18-month-old STING^+/+^ and STING^-/-^ mice lumbar discs for: (A, A’) IL-1β, (B, B’) IL-6, (C, C’) TGF-β), (D, D’) p19, and (E, E’) p21. Images were taken at 10x (scale bar= 100 μm). (n=2-4) discs/animal; 6 animals/genotype, 12-24 total discs/genotype) Dotted lines demarcate different tissue compartments within the disc. Quantitative data represents the mean ± SD. Significance was determined using unpaired t-test or Mann-Whitney test, as appropriate.

